# A complete bacterial genome assembled *de novo* using only nanopore sequencing data

**DOI:** 10.1101/015552

**Authors:** Nicholas J. Loman, Joshua Quick, Jared T. Simpson

## Abstract

A method for *de novo* assembly of data from the Oxford Nanopore MinION instrument is presented which is able to reconstruct the sequence of an entire bacterial chromosome in a single contig. Initially, overlaps between nanopore reads are detected. Reads are then subjected to one or more rounds of error correction by a multiple alignment process employing partial order graphs. After correction, reads are assembled using the Celera assembler. Finally, the assembly is polished using signal-level data from the nanopore employing a novel hidden Markov model. We show that this method is able to assemble nanopore reads from *Escherichia coli* K-12 MG1655 into a single contig of length 4.6Mb permitting a full reconstruction of gene order. The resulting draft assembly has 98.4% nucleotide identity compared to the finished reference genome. After polishing the assembly with our signal-level HMM, the nucleotide identity is improved to 99.4%. We show that MinION sequencing data can be used to reconstruct genomes without the need for a reference sequence or data from other sequencing platforms.

---

The Oxford Nanopore MinION is a portable, single molecule genome sequencing instrument no larger than a typical smartphone. As this instrument directly senses native, individual DNA fragments without the need for amplification, it is able to sequence extremely long fragments (>10 kilobases) of DNA without a reduction in sequence quality ^1^.

The availability of very long reads is important when assembling genomes as they span repetitive elements and anchor repeat copies within uniquely occurring parts of the genome. Many bacterial genomes can be assembled into single contigs if reads greater than eight kilobases are available, as these reads span the conserved ribosomal RNA operon which is typically the longest repeat sequence in a bacterial genome^2^.

The accuracy of the sequence reads is another potentially limiting step for genome assembly; at launch, data from the Pacific Biosciences sequencing instrument was hard to use for *de novo* assembly due to the lack of bioinformatics tools designed for long reads with a high error rate. Several algorithmic improvements have led to this platform becoming the gold standard for genome assembly; Koren *et al.* demonstrated that hybrid techniques could generate contiguous assemblies^3^. This method uses accurate short-read data, or Pacific Biosciences circular consensus reads, to correct error-prone long reads sufficiently for assembly^4^. Non-hybrid *de novo* assemblies of PacBio data in absence of short-read data remained an open problem until Chin *et al.* developed the HGAP (Hierarchical Genome-Assembly Process) assembler^5^. HGAP typically uses a subset of the longest reads in the sequencing data set to use as the input for the assembly process. This subset of long reads is corrected using the entire data set, and this corrected set of reads is assembled with the Celera Assembler. The final assembly is ‘polished’ using a signal-level consensus algorithm to high accuracy. This assembly strategy has revolutionized genome assembly of small genomes and is now being applied to large genomes^6,7^.

Nanopore sequencing data has clear similarities to Pacific Biosciences data, with reads sufficiently long to be of great use in *de novo* assembly^7,8^. Recently, a hybrid approach to assembly which used Illumina short reads to correct nanopore reads prior to assembly was shown to give good results a bacterial and a yeast genome ^9^. However, non-hybrid assemblies have not yet been described. Recent versions of nanopore chemistry (R7.3) coupled with the latest base caller (Metrichor ≥ 1.9) permits read-level accuracies of between 78-85%^1,8^. While this is slightly lower than the latest version of Pacific Biosciences chemistry^6^, we hypothesised that a similar assembly approach where nanopore data reads are self-corrected may result in highly contiguous assemblies.

Overlaps can be detected more easily with higher accuracy data. Therefore for this study we exclusively used high-quality ”passing filter” two-direction (2D) reads. DNA strands which have been read in both directions by the MinION and combined during base-calling are higher quality than the individual template and complement strands ^8^. All 2D reads from four separate MinION runs using R7.3 chemistry were combined. Each run was made using a freshly prepared sequencing library using library protocol SQK-MAP-003 (first run) or SQK-MAP-004 (three further runs). In total, 22,270 2D reads were used comprising 133.6Mb of read data, representing ^~^29x theoretical coverage of the 4.6 megabase *E. coli* K-12 MG1655 reference genome (Supplementary Table 3).

The FASTA sequences for reads were extracted using `poretools`^10^. Potential overlaps between the reads were detected using the `DALIGNER` software^11^. Each read and the reads it overlaps were used as input to the partial order aligner (`POA`) software^12^. `POA` uses a directed acyclic graph to compute a multiple alignment, which we use to determine a consensus sequence. The use of partial order graphs permits a more sensitive reconstruction of consensus sequences in the presence of large numbers of insertions or deletions. This step is analogous to the pre-correction step of the HGAP pipeline, which also uses partial order graphs (implemented in the `pbdagcon` software) to compute a consensus sequence. We run this correction process multiple times, using the corrected reads as the new input. The results of this correction process increased the mean percentage identity from 80.5% to 95.9% after the first iteration and 97.7% on the second iteration (Supplementary Figure 7).

The reads resulting from two rounds of correction were used as input to version 8.2 of the Celera Assembler^13^. The Celera Assembler was run with settings specified in Supplementary Figure 8. The critical parameter for tuning is the overlap minimum identity, which should be double the raw read error rate, in this case set to 6% (`ovlMinIdentity=0.06`). This resulted in a highly contiguous assembly with four contigs, the largest being of length 4.6Mb and covering the entire *E. coli* chromosome (Figure 1). The other three contigs were small (6-15kb) and their sequences were present in the long contig and so we did not include them in further analysis.

**Figure 1:**
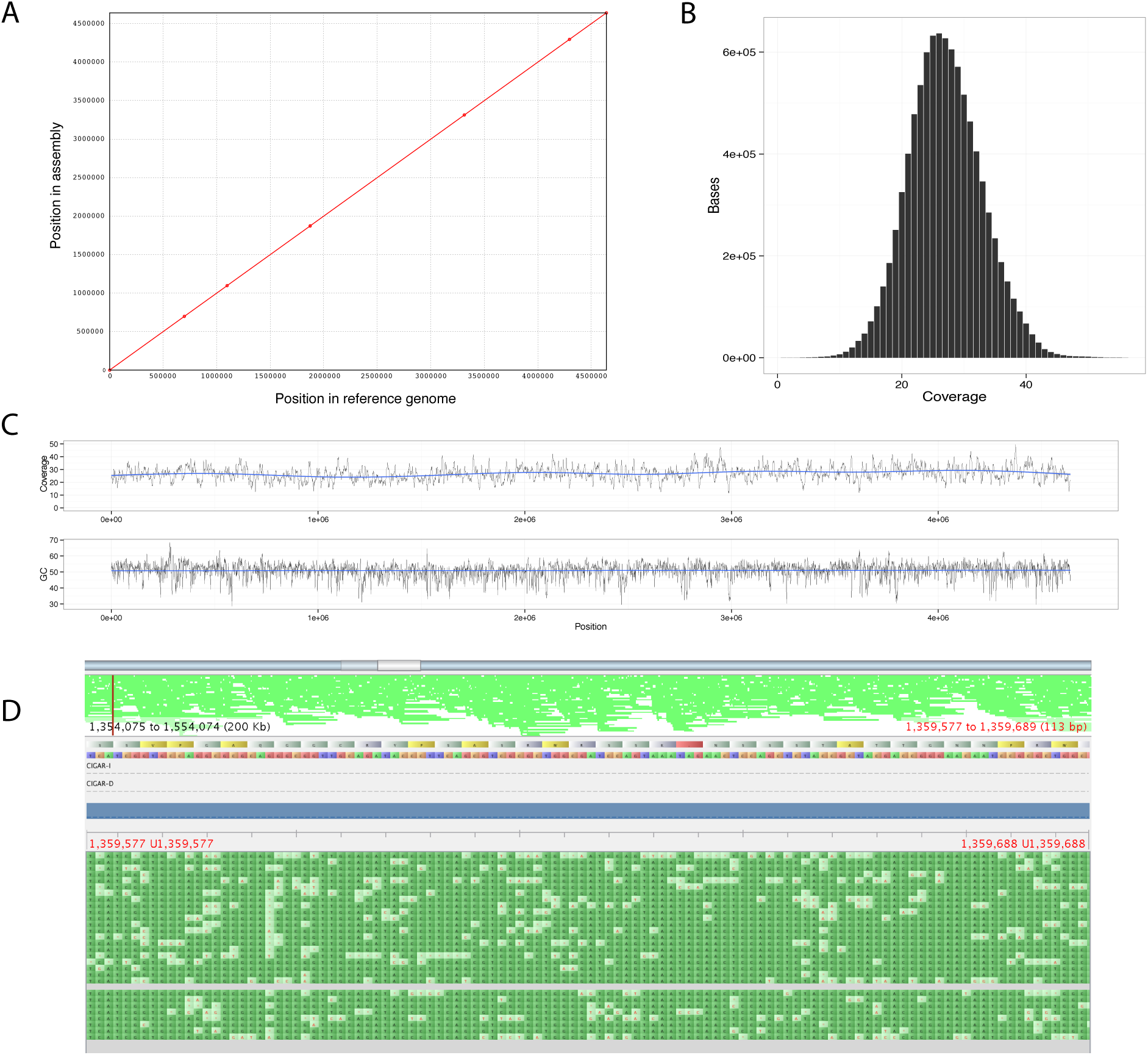
Single contig assembly of *E. coli* K-12 MG1655. Panel A shows a dot-plot comparing the assembly presented here with the reference genome. Panel B shows read coverage as a histogram when nanopore reads are aligned against the assembled genome. Panel C shows read coverage and GC composition across the length of the assembled genome. Panel D shows an illustrative view of the read coverage for a randomly chosen section of the assembly genome and the underlying reads used to construct the assembly as viewed in the genome assembly tool Tablet ^15^.

The initial draft assembly has 3,949 mismatches (85 per 100kb) and 47,395 indels of ≥1 base (1,021 errors per 100kb), as determined by the QUAST validation tool when compared to the *E. coli* K-12 MG1655 reference genome (NC_000913.3)^14^.

We computed a new consensus sequence for the assembly using a hidden Markov model that takes the electric current signals emitted by the MinION as input. This algorithm is described in the supplementary methods. After polishing the assembly using this algorithm the base-level accuracy improved to 99.4%, comprising 1,532 mismatches (33 per 100kb) and 18,387 indels of ≥1 base (396 errors per 100kb).

We explored the sequence context of errors in our assemblies before and after polishing by comparing 5-mer counts between our assemblies and the reference genome. Our initial draft assembly displayed significant under-representation of poly-nucleotide tracts (Figure 2, top panels). This error mode is expected as the MinION relies on a change in electric current to detect base-to-base transitions, which may not occur or may not be detectable as locally repetitive sequence transits the pore. After polishing the assembly the representation of poly-nucleotide tracts improves and the 5-mer counts between the reference genome and the polished assembly are well-correlated (Figure 2, bottom panels).

**Figure 2:**
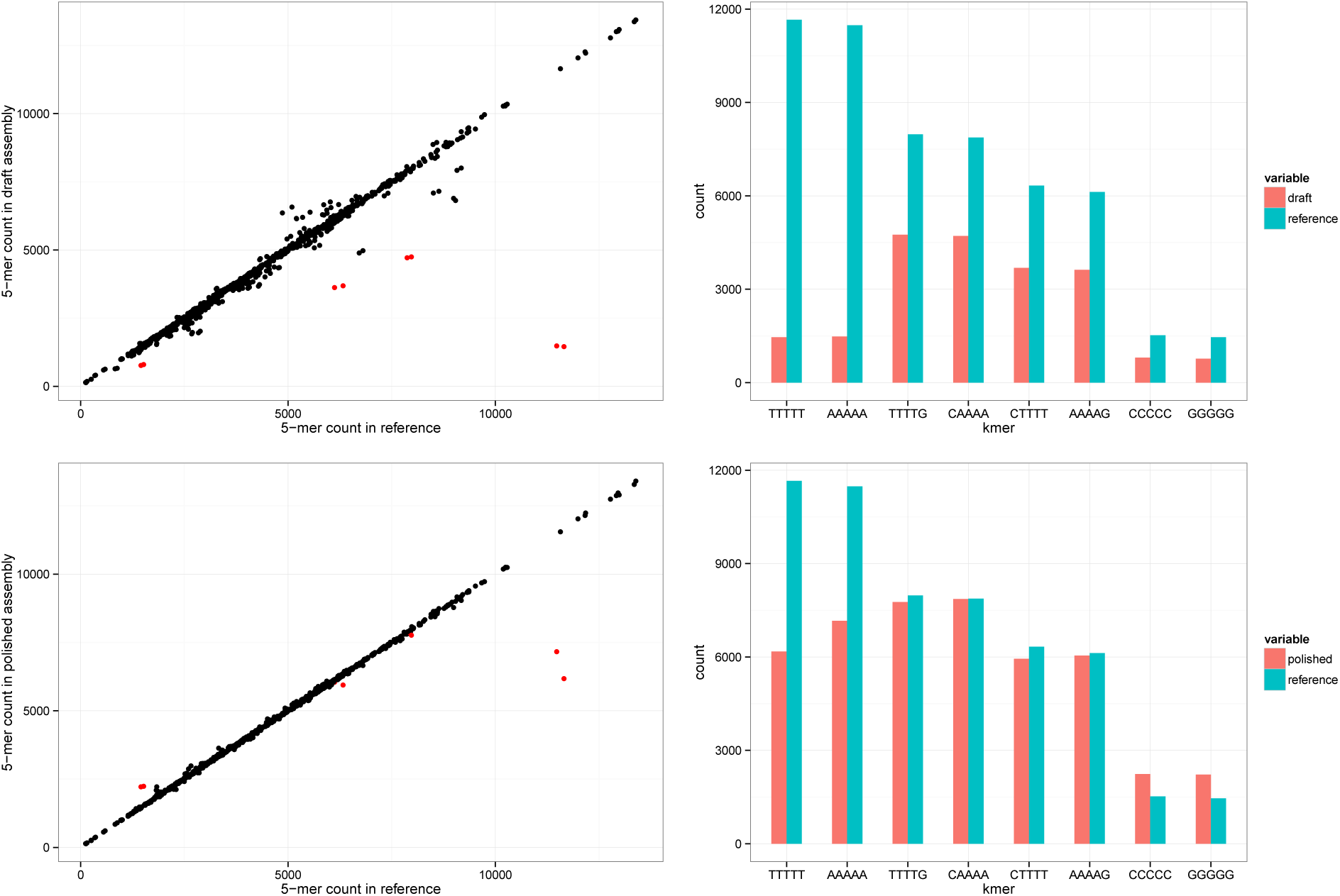
The left panels show the correlation between 5-mer counts in the reference (x-axis) and an assembly (y-axis). The upper left panel is the assembly before signal-level polishing and the lower left panel is after signal-level polishing. The 5-mer counts are well-correlated between the reference and polished assembly (Pearson’s *r* = 0.99). To explore under-representation of 5-mers, we identified sequences where the reference genome had ≥50% more occurrences than the unpolished assembly (red points in the left panels, grouped bars in the right panels). This indicated that the most under-represented 5-mers in our unpolished assembly are those that consist of a single base (TTTTT, AAAAA, CCCCC, GGGGG) or contain a 4-base stretch of a single base (all others, top right panel). After polishing the representation of these difficult motifs is improved (bottom right panel) although `TTTTT` and `AAAAA` remain difficult. In the right panels the bars are ordered by reference occurrence count.

QUAST reported two misassemblies with respect to the reference sequence. On visual inspection of an alignment generated by the Mauve software we determined one large (>500 base) region of difference between our assembly and the reference representing an insertion of a transposase-encoding region into our assembly. This sequence (approximately 750 bases long) is present nine times in the reference genome. Insertion events involving transposons are common in bacterial genomes and can affect wild-type strains commonly used in laboratory experiments, but are difficult to detect using draft assemblies from short-read sequencing technologies.

We conclude that long read data from the Oxford Nanopore MinION can be used to assemble complete bacterial genomes to give an accurate reconstruction of gene order and orientation, without the need to use data from other sequencing technologies. Using signal-level algorithms we are able to improve the per-base accuracy of our assembly further. However, poly-nucleotide tracts remain error-prone. We have not evaluated the impact of increasing sequencing coverage above 30x on accuracy. We expect that improved probabilistic models of the data, possibly incorporating fragment translocation duration information, may result in higher accuracy. The read error correction software, `nanocorrect`, is available at https://github.com/jts/nanocorrect. The signal-level consensus software, `nanopolish` is available at https://github.com/jts/nanopolish. Both programs are freely available under the open-source MIT license.

## Acknowledgements

Data analysis was performed on the Medical Research Council Cloud Infrastructure for Microbial Bioinformatics (CLIMB) cyberinfrastructure. NJL is funded by a Medical Research Council Special Training Fellowship in Biomedical Informatics. JQ is funded by the NIHR Surgical Reconstruction and Microbiology Research Centre. JTS is supported by the Ontario Institute for Cancer Research through funding provided by the Government of Ontario. We are grateful for the staff of Oxford Nanopore for technical help and advice during the MinION Access Programme. We are grateful to the EU COST action ES1103 for funding the authors to attend a hackathon that kickstarted the work presented here. The authors thank Leopold Parts for comments on the manuscript.

## 1 Supplementary Methods

### 1.1 Assembly Pipeline

All the steps described here are additionally documented as an IPython notebook at https://github.com/jts/nanopore-paper-analysis along with the additional scripts to reproduce the analysis presented in this manuscript.

#### Read preprocessing

The `DALIGNER` program was designed for Pacific Biosciences data and expects the input reads to be provided in FASTA format with metadata encoded in the read name. To allow `DALIGNER` to be run on nanopore data, we wrote a program `preprocess.pl` to convert the FASTA output of `poretools` into the format expected by `DALIGNER`.

#### Error Correction

We ran `DALIGNER` on the input reads using default parameters and the procedure described in the documentation to compute read-read overlaps. For each input read, *A,* we queried the `DALIGNER` overlap database to extract the set of reads overlapping *A.* For each read overlapping *A,* we trimmed it to the coordinates matching *A* (as reported by `DALIGNER`) and reverse complemented the sequence if necessary to make the read match the strand of *A.* This set of strand-matched subsequences were written to a FASTA file along with the sequence of *A.* This was the input to `POA`, which was run with parameters `–hb –clustal` using the BLOSUM80 sequence similarity matrix.

The consensus sequence was parsed from the clustal-formatted multiple alignment. We first computed the read depth at each column of the multiple alignment. We then trimmed the multiple alignment by excluding any leading and trailing columns that had depth less than 3. This trimming step is designed to remove uncorrected or unreliable regions of the multiple alignment, which may impact the quality of the downstream assembly. We reported the first consensus record calculated by `POA` (labeled ‘CONSENS0’) over the trimmed region as the consensus sequence for the read.

#### Genome Assembly

After two rounds of error correction we assembled the reads using v8.2 of the Celera Assembly with the parameters file described in Figure 8. The position in the large contig corresponding to the *E. coli* origin of replication was determined through a BLAST homology search for the first 1000 bases of the reference genome, and the contig was circularised using the `minimus2` package in AMOS [12] by the introduction of an artificial contig break. `Minimus2` relies on the `nucmer` component of MUMMER [4] to perform the initial overlap alignments.

### 1.2 Signal-level Consensus Algorithm

The electrical current signal sampled by the MinION inherently contains more information than the base-called reads. We sought to use this information when computing the final consensus sequence of our assembly. In this section we describe our consensus algorithm. We first give a brief overview of the algorithm. We start with our initial draft assembly, *G,* and progressively modify it by making small localized changes. We assess whether the modified sequence, *G*′, increases the probability of the electrical current data for a set of reads. If the probability of the signal data increases we accept *G*′ and iterate. To make these calculations computationally feasible, this process is run over short segments of the genome and seeded by reads aligned to *G.* At the core of this algorithm is calculating the probability of the raw signal data emitted by the MinION given a proposed consensus sequence. We first describe our probabilistic model of the data, then describe how we use this model to compute the consensus sequence for the assembly.

#### Signal-level MinION Data

The MinION continuously samples electrical current flow across the pore. When DNA is detected in the pore, these current samples are turned into *events* by a feature detection algorithm provided by the instrument’s software. The feature detection algorithm is designed to find stepwise changes in the current signal that indicate that the DNA sequence has moved through the pore. For each event the feature detection algorithm records the mean of the current samples (in picoamps), its standard deviation and the total duration of the event (in seconds). Simulated idealized data to illustrate these concepts is shown in Figure 1 and Table 1.

We only use the mean current level in our model but remark that the duration and standard deviation are sources of information that should be considered in future work. When we refer to individual events *e_i_* below we are referring to the mean current level component of the event only. For the purpose of our consensus algorithm, we consider a MinION read to be a sequence of events (*e*_1_*, e*_2_,…, *e_n_*).

**Supplementary Figure 1:**
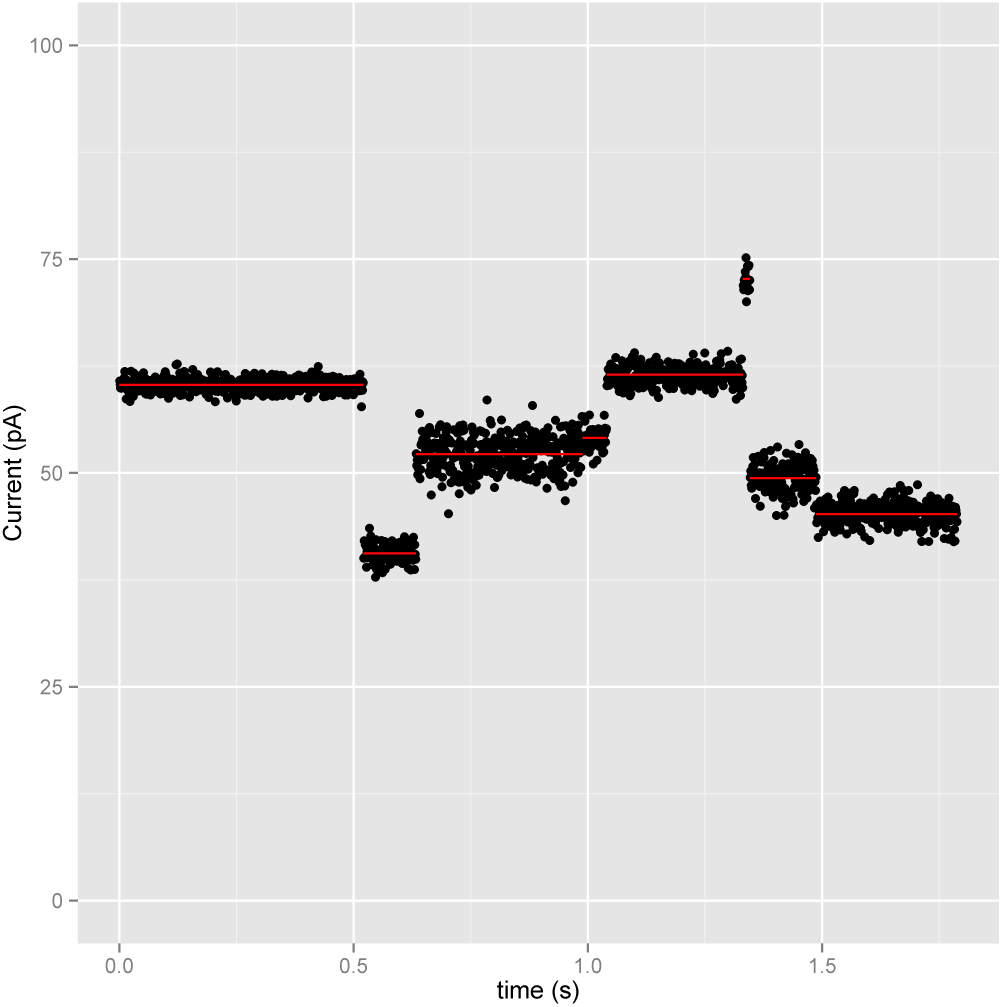
Simulated ideal signal data to illustrate the data that is input into our model. The black points are sampled current levels at a given time. The red lines are the events detected by a feature detection algorithm, which partitions the samples into discrete segments.

#### Modeling Events

The fundamental principle of nanopore-based sequencing is that the amount of current flow through the pore depends on the sequence of DNA that resides in the pore [7, 11]. The MinION model assumes that the current level measured for each event is drawn from a Gaussian distribution with mean and variance dependent on the 5-mer sequence that is in the pore when the samples are taken. Oxford Nanopore provides the parameters for the Gaussians for all 5-mers. We refer to this collection of parameters as **Θ.** The probability of observing event *e_i_* given that 5-mer *k* is the true sequence in the pore is 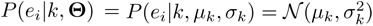.

**Supplementary Table 1:** Events detected from the idealized example shown in Figure 1.

| Event | mean current (pA) | current std dev | duration (s) |
| --- | --- | --- | --- |
| 1 | 60.3 | 0.7 | 0.521 |
| 2 | 40.6 | 1.0 | 0.112 |
| 3 | 52.2 | 2.0 | 0.356 |
| 4 | 54.1 | 1.2 | 0.051 |
| 5 | 61.5 | 1.0 | 0.291 |
| 6 | 72.7 | 1.5 | 0.015 |
| 7 | 49.4 | 1.5 | 0.141 |
| 8 | 45.2 | 1.2 | 0.301 |

#### A Probabilistic Model of MinION Events

We want to calculate the probability of observing an event sequence given a known DNA sequence *S*, *P*(*e*_1_, *e*_2_,…, *e_n_*|*S*, **Θ**). To do this, we represent the known DNA sequence *S* as its constituent 5-mers, *S* = (*s*_1_, *s*_2_,…, *s_m_*). The simplest generative model of the data is where the sequencer moves stepwise from 5-mer to 5-mer, and emits a single event from each 5-mer. This situation is depicted in Figure 2.

**Supplementary Figure 2:**
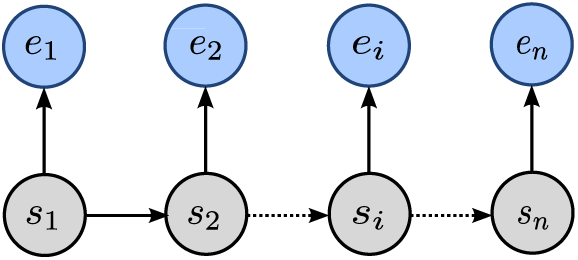
A simple model where each 5-mer (grey circles) emits a single event (blue circles).

In this situation, where there is a perfect one-to-one match between events and 5-mers, calculating the probability of the data is straightforward if we assume independence between events:

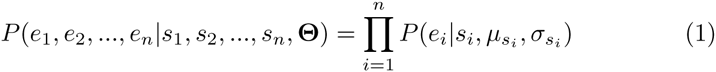

In reality however, we need to account for two types of event detection errors: *skipped* 5-mers, where the system does not detect an event for a 5-mer that transited the pore, and *split events,* where the signal for a single 5-mer is emitted multiple times, presumably because of transient noise that exceeds the event detection threshold.

In this situation, we no longer know which 5-mer emitted each event. To provide motivation for our model we will describe an example from the more complex generative process. Let *S* = (`ACGTA`, `CGTAA`, `GTAAC`, `TAACT`) be the 5-mers of the known DNA sequence. The process starts at the first 5-mer of *S,* `ACGTA`, and emits an event drawn from 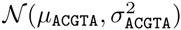. We then choose to stay at `ACGTA`, move to the next 5-mer, `CGTAA`, or skip `CGTAA` and move directly to `GTAAC`. Once a move has been made we can emit another event drawn from the appropriate Gaussian and repeat until we reach the end of the sequence. We can model this process using a hidden Markov model with the structure shown in Figure 3.

**Supplementary Figure 3:**
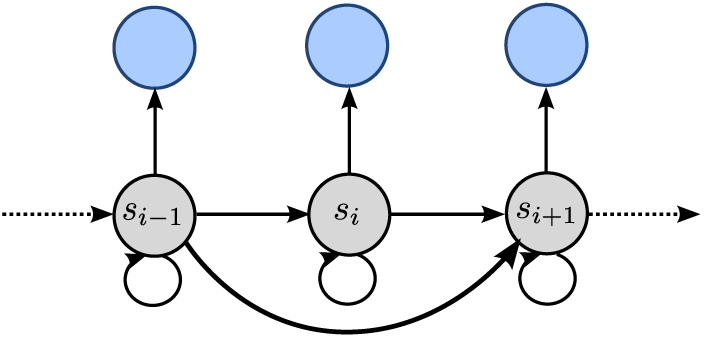
A hidden Markov model allowing self-transitions and jumps over states. This allows us to model split and skipped events.

We will treat the path through the model as the hidden state of the system, *π*, where each entry is a pair *π_i_* = (*i*, *s_j_*) which denotes that at step *i* the system emitted event *i* from 5-mer *s_j_*. We denote the transition probabilities with *P*(*π_i_*|*π_i_*_−1_, *S*) and for convenience define an initial transition *P*(*π*_1_ = (1, *s*_1_)|*π*_0_, *S*) = 1. We can then calculate the joint probability of a path and the data:

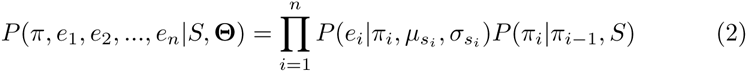

We can calculate the probability of the data given *S* by summing over all possible paths using the Forward algorithm:

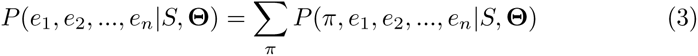

The structure of the model and transition probabilities are implied by the 5-mer sequence *S* as described later.

#### Profile HMM of Events and 5-mer in the Pore

The model drawn above only allows single 5-mers to be skipped. We could add transitions between more distant 5-mers to allow longer jumps but this increases the computational cost to compute *P*(*e*_1_, *e*_2_,…, *e_n_*|*S*, **Θ**) [5]. To allow a similar description of the data in a more computationally convienent structure we can convert this model into a standard Profile HMM. The Profile HMM we use is shown in Figure 4.

The Profile HMM expands the state space to include explicit states for extra events emitted by any 5-mer (*E* states) and silent states that allow 5-mers to be visited without observing an event (*K* states). This model has a repeating block structure with 3 states for every 5-mer in *S.* Each state in a block is indexed by the block’s corresponding 5-mer.

A path *π* through this model consists of states that emit events (incrementing the event index, states *M* and *E*) and states that move from 5-mer to 5-mer (incrementing the 5-mer index, states *M* and *K*). Each entry in the path *π* is again represented as a pair *π_k_* = (*i*, *X*) where *i* is the index of the last event emitted by the path and 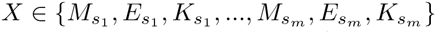. When describing transition probabilities we will use the notation 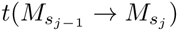 to mean 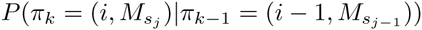 and so on.

To calculate the probability *P*(*e*_1_, *e*_2_,…, *e_n_*|*S*, **Θ**) we again use the Forward algorithm which fills in a dynamic programming matrix of size 3*mn* to sum over all paths through the model. We refer the reader to section 5.4 of Durbin et al. [5] for the recursive definition of the dynamic programming matrix for a Profile HMM.

**Supplementary Figure 4:**
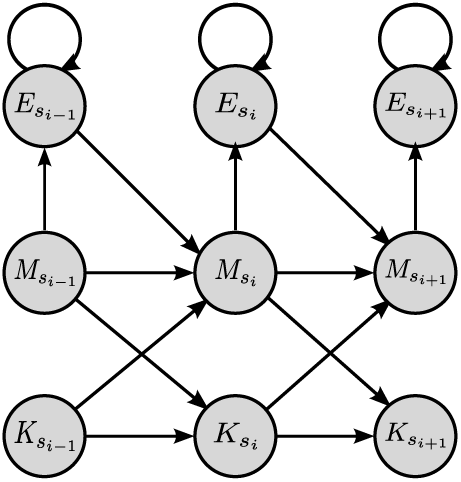
The state structure of the Profile Hidden Markov Model we use to calculate the probability of a sequence of events given a known sequence.

#### Transition Probabilities

The transition probabilities of the model are crucial to capturing the behaviour of the pore. Most importantly, we must accurately model the probability of not observing a signal for a particular 5-mer. If the signals for two adjacent 5-mers, *s_i_*_−1_ and *s_i_*, are similar the feature detection algorithm may not register a change in current. To make this explicit, consider the situation where the sequenced DNA is GAAAAAAC. There is no chance to register the `AAAAA` → `AAAAA` transition as the current signals for these 5-mers are identical by definition. Therefore the probability of an event sequence generated by *S* given two candidate sequences *S*_1_ = GAAAAAAC and *S*_2_ = `GAAAAAC`, which only differ by the undetectable length of the poly-A run, should be very similar, *P*(*e*_1_, *e*_2_,…, *e_n_*|*S*_1_, **Θ**) ≈ *P*(*e*_1_, *e*_2_,…, *e_n_*|*S*_2_, **Θ**).

We model this by making the transition probability to *K* states a function of the absolute difference between expected current for the adjacent 5-mers 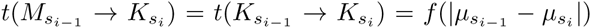. We use a lookup table for *f* that was calculated from a sample of reads matched to an earlier draft assembly. The lookup table stores probabilities in bins of 0.5 pA.

While the majority of skipped 5-mers are due to signal similarity, we also observe skips of 5-mers whose signal differs substantially from the preceding 5-mer. These occurrences are presumably due to events that were too short to reliably detect and hence discarded by the feature detection algorithm.

The probability of transitioning to an *E* state is independent of the 5-mer. Similar to the *f* table, we learned the probability of emitting an additional event *p_me_* from a sample of reads. We calculate 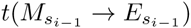 using *p_me_* and the probability of not transitioning to the *K* state:

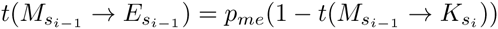

We also estimated the probability of staying in the *E* state, *p_ee_*, from the same sample of reads and set 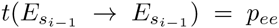. Later we describe how these transition probabilities are trained to capture the characteristics of individual MinION reads. The complete set of transition probabilities for our model is as follows:

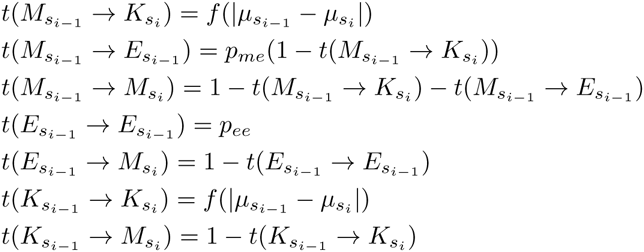

#### Emission Distributions

To complete the model we need emission distributions for each state. The emission distribution for the *M* state are the Gaussians described above. The emission distribution for the *E* states are nearly the same however we noted higher variance for the duplicated events so we scale the standard deviation by a factor of *v* = 1.75. As described above, no events are emitted from the *K* states.

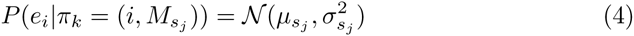

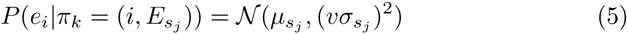

This completes the description of our HMM.

**Supplementary Table 2:**
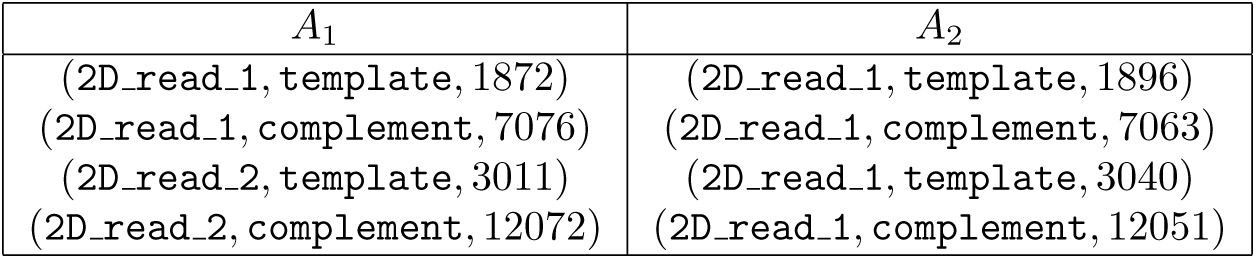
An example of the first two reads for the first two anchors shown in Figure 5.

| $A_1$ | $A_2$ |
| --- | --- |
| (2D_read_1, template, 1872) | (2D_read_1, template, 1896) |
| (2D_read_1, complement, 7076) | (2D_read_1, complement, 7063) |
| (2D_read_2, template, 3011) | (2D_read_1, template, 3040) |
| (2D_read_2, complement, 12072) | (2D_read_1, complement, 12051) |

### 1.3 Computing the Consensus Sequence

Our consensus pipeline begins by mapping the uncorrected 2D MinION reads to the draft assembly, *G*. To do this we use `bwa mem` with option `-x ont2d` [8]. After the read mapping step, we segment the genome into 10 kilobase segments which overlap by 200bp. For each 10kb segment we extract the subsequence of each read that maps to the segment and create a multiple alignment using POA as in our error correction algorithm. The multiple alignment for the segment is the input into our consensus algorithm.

The first row of the multiple alignment is the sequence of the draft assembly for the segment and the remaining rows are the 2D reads mapped to the segment. We iterate over the columns of the multiple alignment and select an *anchor* every 50bp of the draft assembly’s sequence. These anchor points define a mapping between bases in the draft assembly and the current signal events for each strand (template and complement) of the aligned 2D MinION reads. Later, when evaluating the probability of the data in our HMM we treat each strand of a 2D read independently. An anchor *A_i_* is a set of tuples specifying a MinION read identifier, a strand identifier and the index of the nearest event for the base aligned to the draft assembly. We use the anchor positions to extract subsequences of the draft assembly, the 2D MinION reads and their template and complement events to input into the HMM. An example is in Figure 5 and Table 2.

**Supplementary Figure 5:**
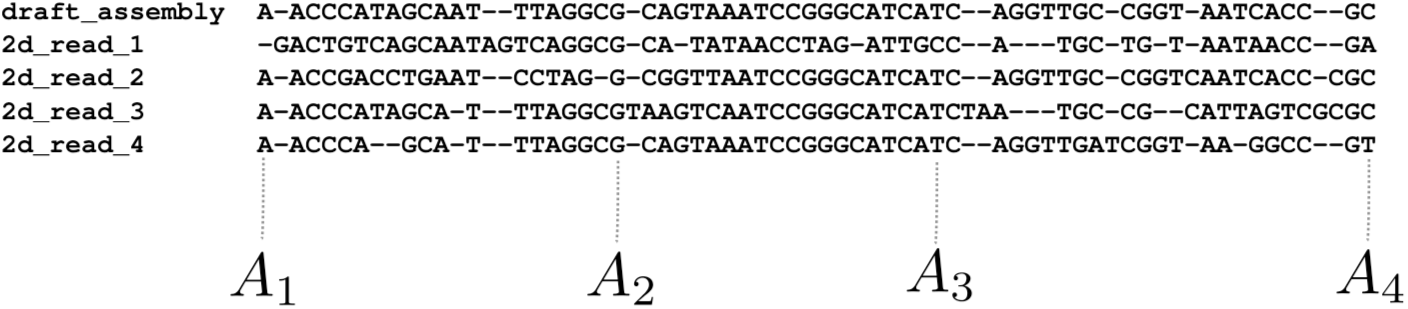
Anchor positions are selected from the multiple alignment which store mappings between bases and MinION events for every 2D read mapped to a segment. In this example, anchors are selected every 20bp along the draft assembly row.

#### Training the HMM Transitions

Prior to computing the consenseus sequence of a segment, we adjust the transition probabilities for both strands of the MinION reads. We do this by aligning the events for each read strand to the draft assembly sequence using the Viterbi algorithm, guided by the anchor positions. For each read, we count the number of transitions between the states of the Profile HMM and update the *f* table, *p_me_* and *p_ee_* accordingly. As a read may not have have enough events to estimate the *f* table across the full range of values we use pseudocounts from the original *f* table to initialize this update.

#### Consensus Algorithm

The consensus algorithm recomputes the sequence of the draft assembly between anchors *A_i_* and *A_i_*_+2_. After the new consensus sequence is computed, we update the sequence of the assembly and the event-to-assembly mappings for *A_i_*_+1_. This update step uses the Viterbi algorithm to align events to the new consensus sequence. The purpose of this pattern - jumping over an anchor and realigning - is to progressively improve the quality of the event-to-assembly alignments recorded in the anchors.

Let 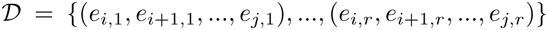 be the set of event subsequences for the *r* strands of the 2D MinION reads that are anchored at both *A_i_* and *A_i_*_+2_. Our goal is to compute a consensus sequence *C* that maximizes the probability of the data 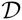. We do this by initializing *C* to the sequence of the draft assembly between the two anchors. We then propose a set of candidate sequences 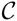 which are derived from the current consensus *C*. We use our HMM to select the sequence that maximizes the probability of the data:

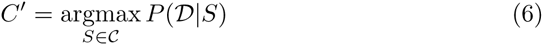

where

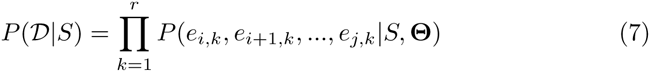

The current consensus sequence is always included in the set of candidates. If the current consensus sequence is selected by equation 6, the process stops.

The set of all possible candidate sequences between *A_i_* and *A_i_*_+2_ is too large to test every possibility. Instead, we use two methods to propose small sets of candidate sequences derived from the current consensus sequence *C*. The first method makes edits to *C* based on sampling alternative substrings from the reads. The second method, inspired by the Quiver algorithm [1], generates all strings within edit distance of one of *C*. This pair of methods, the first allowing large changes that must be observed by at least one read, the second exhaustively trying all one base edits, balances between compute time and thoroughly exploring the space of candidate consensus sequences.

The first method, which we call the `block replacement` algorithm, computes the longest common subsequence of 5-mers between *C* and an aligned 2D read, *R*. This algorithm partitions *C* and *R* into contiguous regions where they share matching 5-mers and regions where they do not. For each pair of consecutive matching regions, we replace the sequence in *C* with the sequence in *R* and add this derived sequence to the set 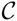. This algorithm generates a new candidate sequence for each pair of matching regions between *C* and *R* and is run for all 2D reads that are aligned between the anchors.

The second method, which we call the mutation algorithm, simply makes all one base substitutions, one base insertions and one base deletions to *C* to generate the candidate set 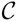.

These algorithms are run iteratively. We initialize the current consensus *C* to the draft genome sequence between the pair of anchors. We then generate candidate set 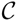 using the `block replacement` algorithm. We select the best sequence *C*′ using equation 6. If *C*′ is not *C*, we set *C* ← *C*′ and repeat the process. Once no more improvements to the consensus sequence are made, we run this procedure again except using the `mutation` algorithm to generate 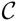. Once this procedure converges we update the assembly to contain the final consensus sequence. We then update the event alignments for *A_i_*_+1_ and move on to the next pair of anchors. This process continues until the entire 10kbp sequence has been processed. Once all 10kbp segments have been processed, which occurs in parallel, we merge the segments together at the overlapping 200bp ends.

#### Culling Unlikely Candidate Sequences

While we restrict our candidate sequences to those that are close to the current consensus *C*, we still generate many sequence that are unlikely to be the true sequence of the genome between a pair of anchors. To avoid spending computation time fully evaluating equation 6 for these sequences we periodically cull unlikely sequences from 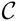. We do this by progressively evaluating equation 6 using subsets of *k* elements of the full data 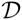. We will refer to these subsets as 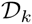.

To retain a sequence *S* in 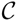, we require that either *k*/5 event sequences in 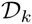 have greater probability given *S* than the current consensus sequence *C* or that 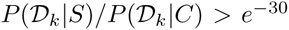. If both of these conditions fail, we remove *S* from 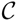.

These checks are performed after processing 5 event sequences, eg for subsets 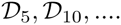. The conservative threshold that we set on the ratio between probabilities is to avoid culling the true sequence from 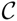 at the cost of retaining some unlikely sequences.

#### Event Preprocessing

The events recorded for a read may differ from the reference pore model that Oxford Nanopore has provided. To account for this the MinION’s software calculates parameters to globally shift and scale the mean and variance of the provided 5-mer model to better match the observed data. Additionally a correction is applied to the event signals to correct for a tendency of the signal to drift over time. We apply these corrections, shifts and scales to the pore model and events prior to input into our HMM.

#### Read Filtering

During development we noticed that some MinION reads contained observations that were outside of the expected range of signals. To discard these reads rather than having the erroneous observations dominate the probability in equation 6, we filtered the input events by computing the probability of the data given the original draft reference sequence and discarding any strands whose log-scaled probability divided by the number of events is greater than 3.5. Better quality control of the input data is an area of future improvement.

#### Source Code and Commands Run

The source code for our consensus algorithm is freely available and open source under the MIT license at https://github.com/jts/nanopolish. A description of how we generated the consensus sequence for the assembly in this paper is documented on github:

https://github.com/jts/nanopore–paper–analysis/blob/master/polishing_commands.txt

### 1.4 Assembly Analysis

#### Analysis of read accuracy

For the analysis of nanopore sequence read accuracy, reads were mapped to the *E. coli* K-12 MG1655 reference genome using `BWA-MEM 0.7.10-r960-dirty` [8]. The resulting BAM files were processed with the `expand-cigar.py` script and mismatches, insertions and deletions counted using the `count-errors.py` script from Aaron Quinlan’s `nanopore-scripts` package (https://github.com/arq5x/nanopore-scripts).

#### Dotplot analysis

The circularised contig was analysed through the `nucmer` component of MUMMER [10] and the delta file visualised using `mummerplot`.

#### Coverage analysis

The sorted, aligned BAM file of raw reads against the assembly was analysed using the `genomeCoverageBed` module of `bedtools` [10]. Per-base coverage was averaged over 1000 base segments and plotted in R using `ggplot2`.

#### GC analysis

The GC plot was constructed from the assembly using a Python script using Biopython averaging GC-content over 1000 base segments [2].

#### QUAST Analysis

The circularised contig, both before and after polishing was analysed by QUAST 2.3 [6] which was run with default parameters and supplying the `NC_000913.3` reference genome as the comparator.

#### Mauve Analysis

The circularised contig was analysed by running the `progressiveMauve` package on the command-line with option `–collinear` and visualised in the Mauve Genome Viewer. [3].

#### 5-mer Analysis

To explore over and under-representation of 5-mers, we wrote a program to count the number of occurrences of the 1024 different 5-mers in a set of sequences. This program counts 5-mers on the forward and reverse strands independently. The analysis used the counts for the forward strand of the reference and the forward strand of the largest contig of our assemblies. We plotted the results using `R` and the `ggplot2` library.

## 2 Supplementary Figures and Tables

**Supplementary Figure 6:**
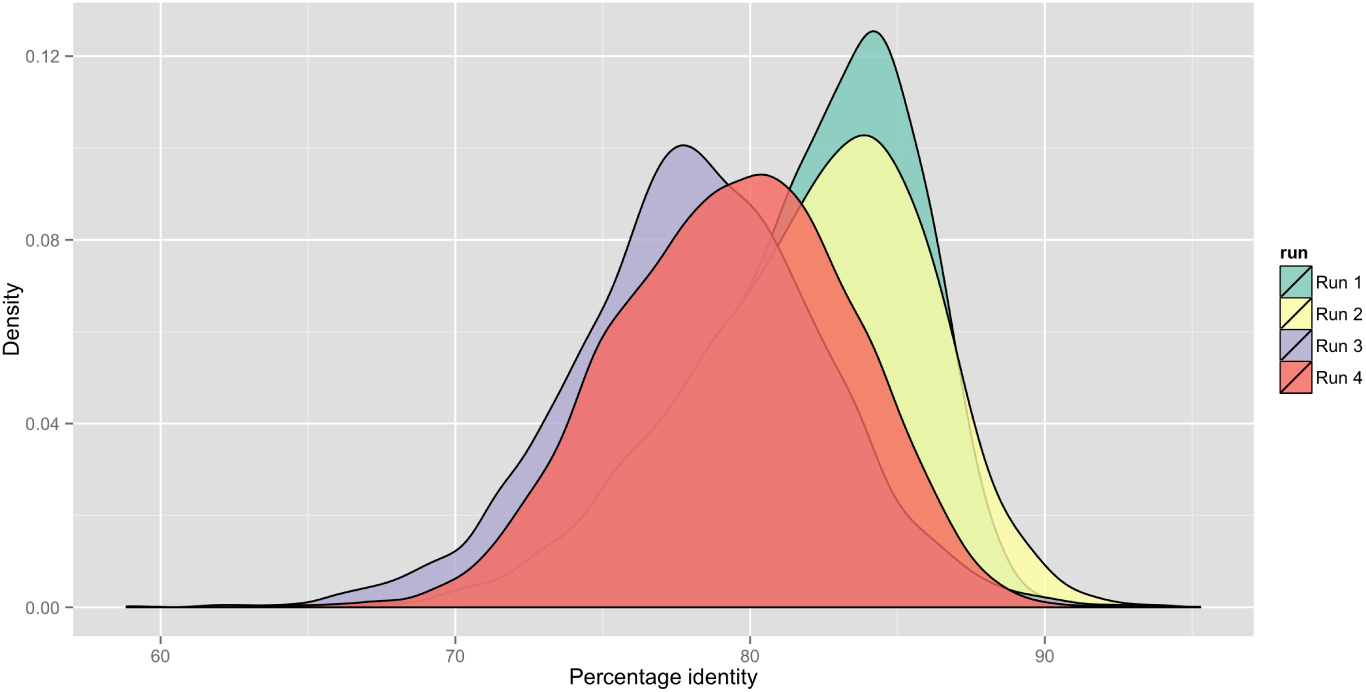
Kernel density plot showing the accuracy of reads from the four individual MinION runs used to generate the *de novo* assembly. The mean accuracy varies from 78.2% (run 3) to 82.2% (run 1).

**Supplementary Figure 7:**
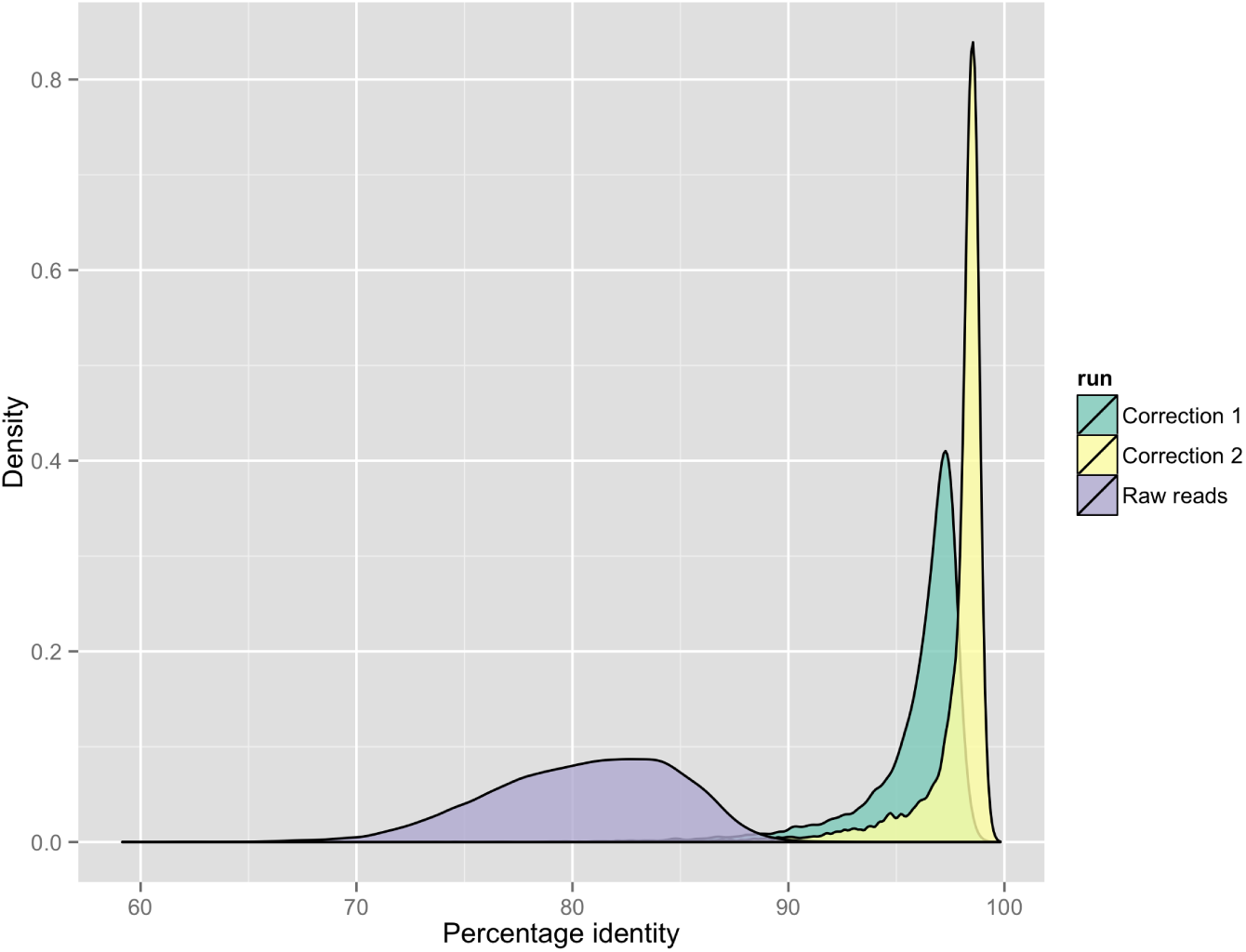
Kernel density plot demonstrating the raw nanopore read accuracy and effect of two rounds of error correction on accuracy. The mauve area represents uncorrected sequencing reads, where the green area shows the improvement in accuracy after the first round of correction and the yellow shows improvement from the second round of correction. Further rounds of correction did not improve the accuracy further.

**Supplementary Figure 8:**
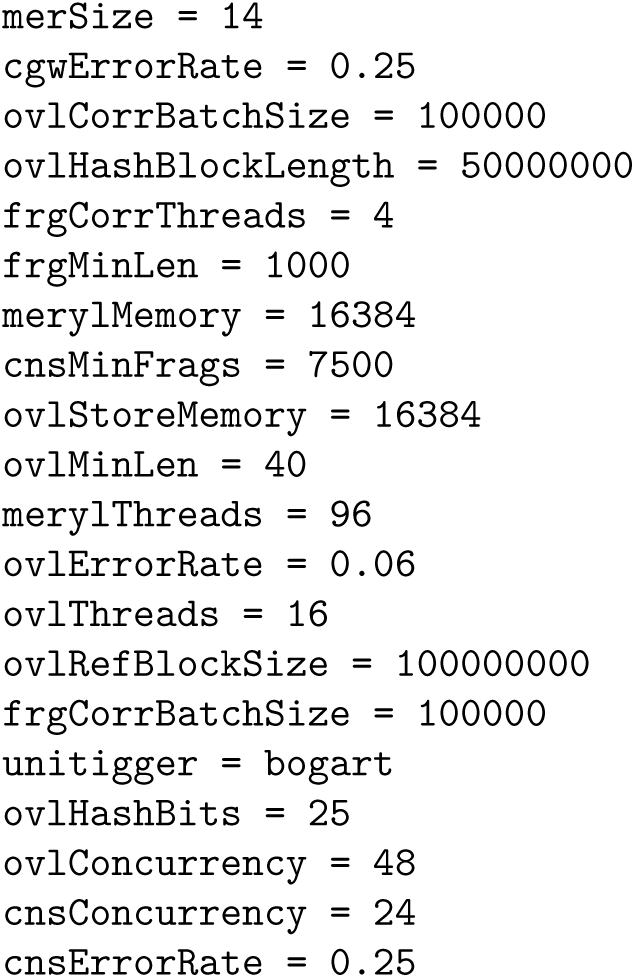
Spec file for Celera Assembler.

**Supplementary Table 3:** Oxford Nanopore MinION datasets used in this study.

| Description | Pass 2D | Mean length | Accession |
| --- | --- | --- | --- |
| R7.3 chemistry, Metrichor 1.9, SQK-MAP-003 | 8451 | 6183 | ERX708228 [9] |
| R7.3 chemistry, Metrichor 1.9, SQK-MAP-004 | 2855 | 5604 | ERX708229 |
| R7.3 chemistry, Metrichor 1.9, SQK-MAP-004 | 5639 | 3885 | ERX708230 |
| R7.3 chemistry, Metrichor 1.9, SQK-MAP-004 | 5325 | 8158 | ERX708231 |

**Supplementary Table 4:** Results from QUAST when comparing the draft circular contig (”circular”) and the polished circular contig (”circular_polished”) against the *E. coli* K12 MG1655 reference genome (accession `NC_000913.3`).

| Assembly | circular | circular_polished |
| --- | --- | --- |
| # contigs ($\geq 0$ bp) | 1 | 1 |
| # contigs ($\geq 1000$ bp) | 1 | 1 |
| Total length ($\geq 0$ bp) | 4593653 | **4639191** |
| Total length ($\geq 1000$ bp) | 4593653 | **4639191** |
| # contigs | 1 | 1 |
| Largest contig | 4593653 | **4639191** |
| Total length | 4593653 | **4639191** |
| Reference length | 4641652 | 4641652 |
| GC (%) | 51.07 | 50.94 |
| Reference GC (%) | 50.79 | 50.79 |
| N50 | 4593653 | **4639191** |
| NG50 | 4593653 | **4639191** |
| N75 | 4593653 | **4639191** |
| NG75 | 4593653 | **4639191** |
| L50 | 1 | 1 |
| LG50 | 1 | 1 |
| L75 | 1 | 1 |
| LG75 | 1 | 1 |
| # misassemblies | 2 | 2 |
| # misassembled contigs | 1 | 1 |
| Misassembled contigs length | **4593653** | 4639191 |
| # local misassemblies | **2** | 4 |
| # unaligned contigs | 0 + 0 part | 0 + 0 part |
| Unaligned length | 0 | 0 |
| Genome fraction (%) | **100.000** | 99.995 |
| Duplication ratio | **0.990** | 1.000 |
| # N's per 100 kbp | 0.30 | **0.00** |
| # mismatches per 100 kbp | 85.08 | **33.01** |
| # indels per 100 kbp | 1021.08 | **396.15** |
| Largest alignment | 2739950 | **2766640** |
| NA50 | 2739950 | **2766640** |
| NGA50 | 2739950 | **2766640** |
| NA75 | 1852939 | **1871782** |
| NGA75 | 1852939 | **1871782** |
| LA50 | 1 | 1 |
| LGA50 | 1 | 1 |
| LA75 | 2 | 2 |
| LGA75 | 2 | 2 |

